# Systematic variation in the temperature dependence of bacterial carbon use efficiency

**DOI:** 10.1101/2020.09.14.296095

**Authors:** Thomas P. Smith, Tom Clegg, Thomas Bell, Samrāt Pawar

## Abstract

Understanding the temperature dependence of carbon use efficiency (CUE) is critical for understanding microbial physiology, population dynamics, and community-level responses to changing environmental temperatures ^1,2^. Currently, microbial CUE is widely assumed to decrease with temperature ^3,4^. However, this assumption is based largely on community-level data, which are influenced by many confounding factors ^5^, with little empirical evidence at the level of individual strains. Here, we experimentally characterise the CUE thermal response for a diverse set of environmental bacterial isolates. We find that contrary to current thinking, bacterial CUE typically responds either positively to temperature, or has no discernible temperature response, within biologically meaningful temperature ranges. Using a global data-synthesis, we show that our empirical results are generalisable across a much wider diversity of bacteria than have previously been tested. This systematic variation in the thermal responses of bacterial CUE stems from the fact that relative to respiration rates, bacterial population growth rates typically respond more strongly to temperature, and are also subject to weaker evolutionary constraints. Our results provide fundamental new insights into microbial physiology, and a basis for more accurately modelling the effects of shorter-term thermal fluctuations as well as longer-term climatic warming on microbial communities.

---

The efficiency with which bacterial populations convert organic carbon into biomass, generally termed Carbon Use Efficiency (CUE), is a key physiological measure that ultimately determines the rate at which whole microbial communities decompose organic matter and release CO_2_ ^1^. Therefore, CUE is a key parameter in global carbon cycle models ^2,6^, as well as models of soil biogeochemical processes ^3,7,8^ and marine particle export ^9^. CUE is typically quantified as the ratio of carbon allocated to biomass production relative to the total carbon assimilated ^1,10^:

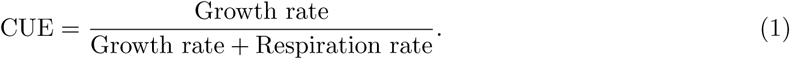

The denominator of this quantity is the sum of rates of carbon allocation to growth and respiration, a common approximation where direct measurements of uptake are not feasible ^1,11,12,13^. High CUE values imply increased biomass production (sequestration) relative to CO_2_ release due to respiration, and vice versa ^1^. Microbial CUE varies with environmental conditions such as resource stoichiometry and availability ^11^, and physical parameters such as pH and temperature ^13,14^. CUE values reported from environmental samples are therefore generally much lower than may be expected from theoretical calculations ^10^, as microbial communities are very rarely operating under conditions for optimal growth efficiency. The response of microbial CUE to changes in environmental temperature is particularly important, both for understanding how microbial communities respond to spatial and temporal variation in temperature, as well as for predicting the effects of climate change on carbon cycling.

Currently, models of organic matter decomposition typically assume a decrease in microbial CUE with temperature ^3,4,15^. This is based on the premise that microbial respiration rate displays a stronger thermal sensitivity than growth rate ^1,4^, implying that growth efficiency declines with temperature. However, results from empirical studies in both soil ^16,17^ and aquatic systems ^1,14,18^ are ambiguous, with studies variously finding decreases ^7^, increases ^12^, or little to no change in CUE with temperature ^19,20,21^. Recent work at the level of single bacterial strains has also challenged this generalisation, finding variable CUE thermal responses between taxa ^13^. However, most previous studies have focused on the CUE of whole microbial consortia in environmental samples, permitting limited mechanistic understanding of these responses. This is because temperature-driven community composition changes are expected to influence CUE ^22^, and also because it is difficult to control for temperature-driven changes in nutrient availability in the medium ^5,19^. This uncertainty about strain-level thermal responses of bacterial CUE severely limits our ability to understand responses of microbial populations to warming, and build mechanistic models of community-level responses.

Here, we quantify CUE using laboratory experiments at the level of single strains for 29 aerobic environmental bacterial isolates spanning 9 families within 3 phyla. We combine this with a data-synthesis of > 400 growth and respiration thermal performance curves spanning most major culturable bacterial phyla ^23^, to uncover general patterns in the temperature-dependence of CUE.

We first made precise the relationship between the thermal performance curve (TPC) of CUE and that of its underlying metabolic traits using a mathematical model (Methods). This model allows us to express the thermal sensitivity of CUE (its apparent activation energy, *E*_CUE_) as a function of the sensitivities of growth rate (*µ*) and respiration rate (*R*) (as activation energies *E*_*µ*_ and *E*_*R*_, respectively) within the population’s Operational Temperature Range (OTR) (Fig. 1A). *E*_CUE_ therefore describes a population’s change in CUE with temperature across the OTR. Specifically, within the OTR, CUE decreases with temperature (negative *E*_CUE_) if the thermal sensitivity of respiration is greater than that of growth, and vice versa.

**Figure 1:**
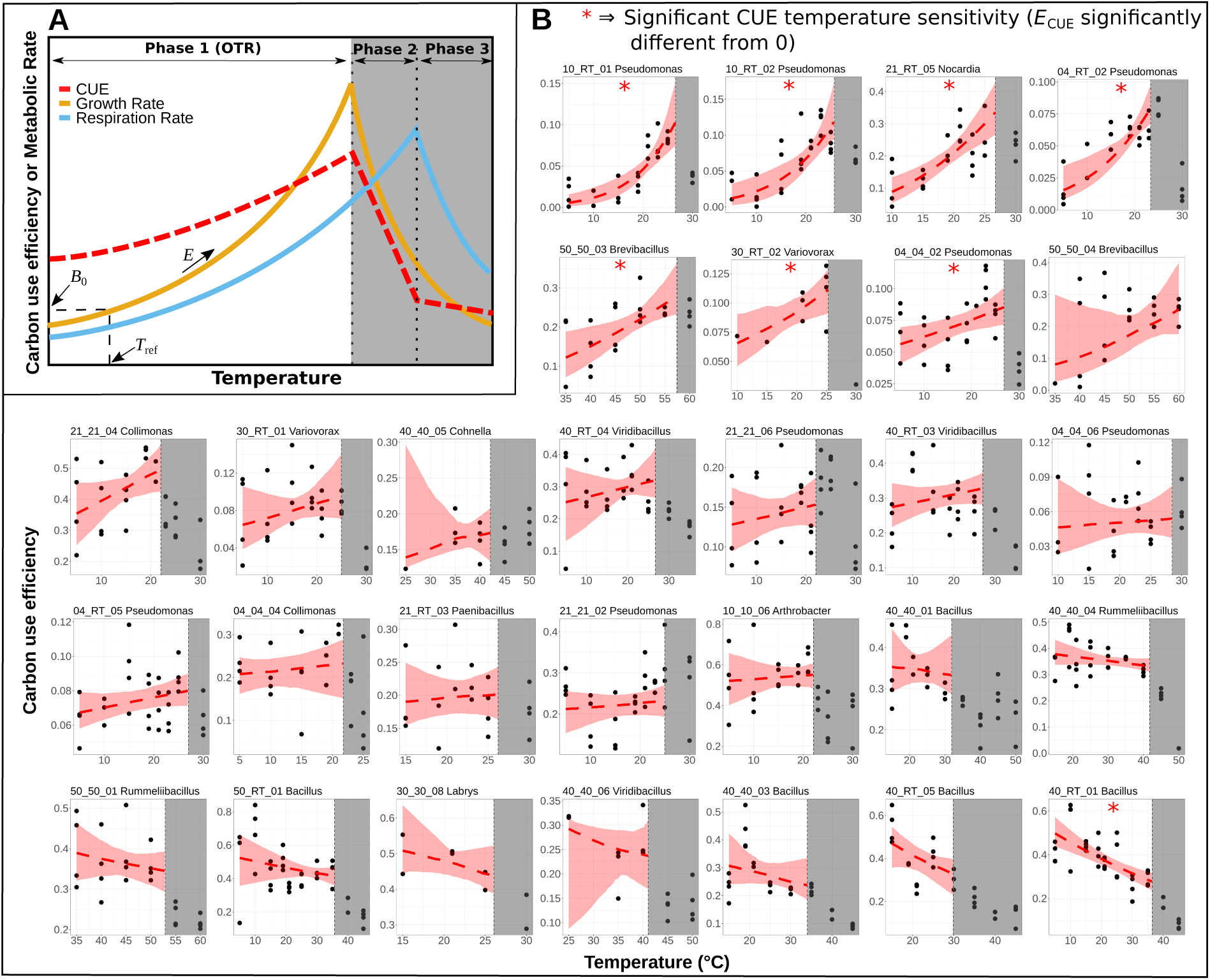
The temperature dependence of carbon use efficiency. **A**. Growth (orange) and respiration (blue) show a unimodal thermal performance curve (TPC) with temperature. The portion of the TPC within the population’s operational temperature range (OTR)—the unshaded region—can be modelled using the Boltzmann-Arrhenius (BA) equation (eq. 4 in Methods; model parameters labeled on growth rate TPC). The upper limit of the OTR is defined by *T*_pk,*µ*_, the temperature at which growth rate peaks. The difference in BA equation parameters between growth and respiration determines the TPC of CUE (red dashed line). **B**. The TPCs of the within-OTR CUE for each of 29 bacterial strains (up to 4 replicates at each temperature). The header for each plot gives the strain ID code (Supplementary Table S1) and the bacterial genus. The red dashed line is the TPC of CUE within the OTR, calculated as the median of the responses of 1000 bootstrapped fits of the TPCs of *µ* and *R* to the Boltzmann-Arrhenius model (Methods). The red shaded area is the (bootstrapped) 95% confidence envelope around the CUE TPC.

We then estimated *E*_*R*_ and *E*_*µ*_ for each of the 29 bacterial strains by fitting the thermal response of growth and respiration rate within its OTR to the Boltzmann-Arrhenius TPC model (eq. 4) (Methods). To characterise the TPCs of the two traits, we measured growth and respiration rates at the same time-point of (exponential) population growth, over the same timescale, overcoming a key limitation of many previous such studies (Methods). Across our dataset, we find that the majority of strains (21/29) display a non-significant response of CUE to temperature within their OTR (Figs. 1B & 2). Seven strains show a significant increase in CUE with temperature, while only one strain shows a significant decrease in CUE with temperature (Figs. 1B & 2, Supplementary Table S2). Furthermore, we find that strains showing a positive CUE thermal response tend to be those with lower CUE in general, whilst the opposite is true for high efficiency strains (linear regression, intercept = 0.44, slope = −0.76, *F*_1,27_ = 10.86, *p* = 0.0028, Fig. 2B). Although by eye there appears to be some curvature, a straight line is preferred by AIC over a polynomial using linear regression. These responses are taxonomically structured, with lower efficiency *Proteobacteria* showing positive temperature responses and higher efficiency *Firmicutes* tending towards negative CUE thermal responses. Also, although the thermal optima for growth (*T*_pk,*µ*_) and respiration (*T*_pk,*R*_) are highly correlated (Pearson’s *r* = 0.91), growth rates generally peak at lower temperatures than respiration rates (*T*_pk,*µ*_ < *T*_pk,*R*_, paired *t*_23_ = 4.996, *p* < 0.001 Fig. 3A). This validates our assumption of a monotonic CUE thermal response within the OTR (Fig. 1).

**Figure 2:**
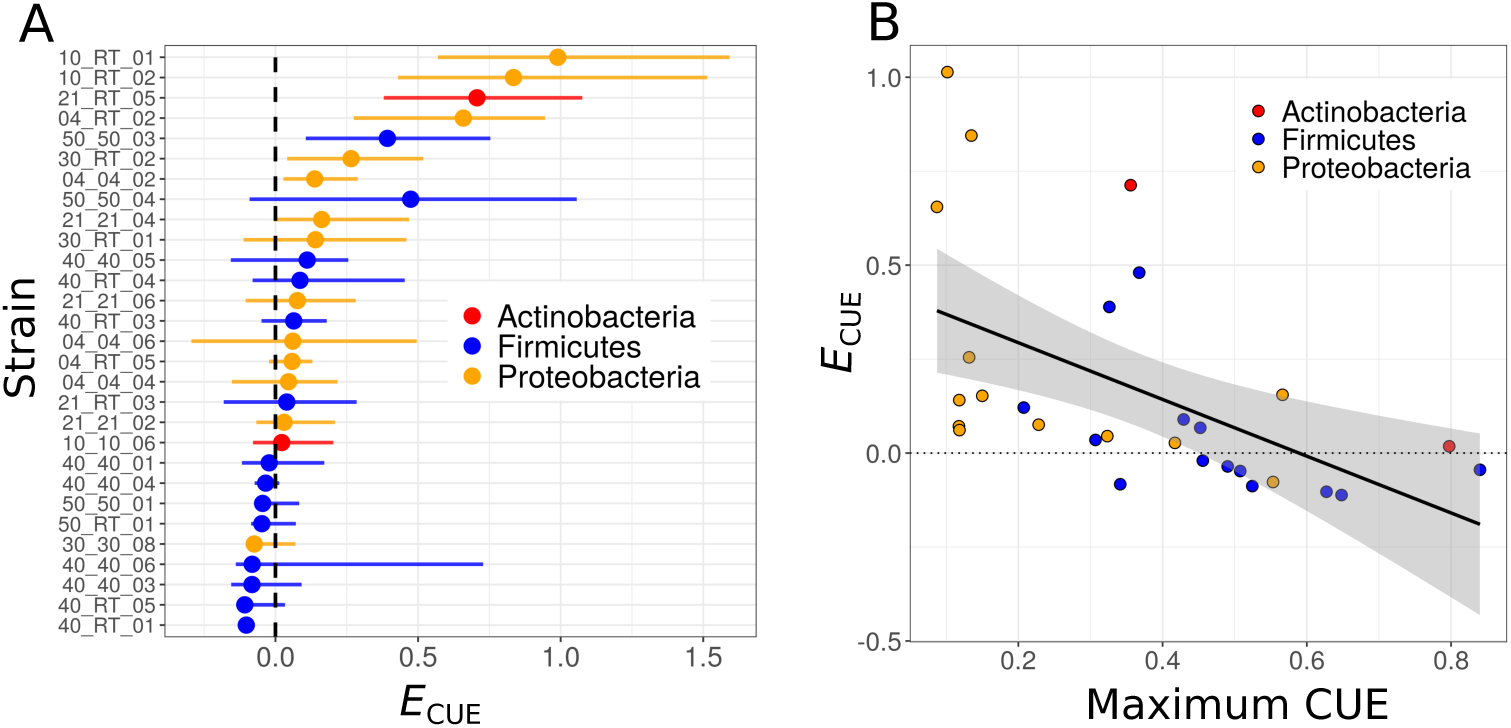
The thermal sensitivity of CUE varies across bacterial taxa. **A** Median bootstrapped *E*_CUE_ with 95% confidence intervals (CIs), strains ordered by the directionality of their response, from positive to negative, and coloured by phylum. Seven strains have a lower CI that falls above zero (black, dashed line), indicating a positive CUE thermal sensitivity within the OTR. The majority of strains’ CIs include zero, indicating insignificant directionality (CUE TPC is thermally insensitive). A single strain “40_RT_01” displayed a significantly negative thermal response for CUE. **B** There is a significant negative relationship (linear regression *p* = 0.00275, black line with grey confidence envelope) between the measured CUE for each strain, and its CUE thermal sensitivity (*E*_CUE_), *i.e*., less efficient strains are able to increase their efficiency with temperature, while high efficiency strains cannot.

**Figure 3:**
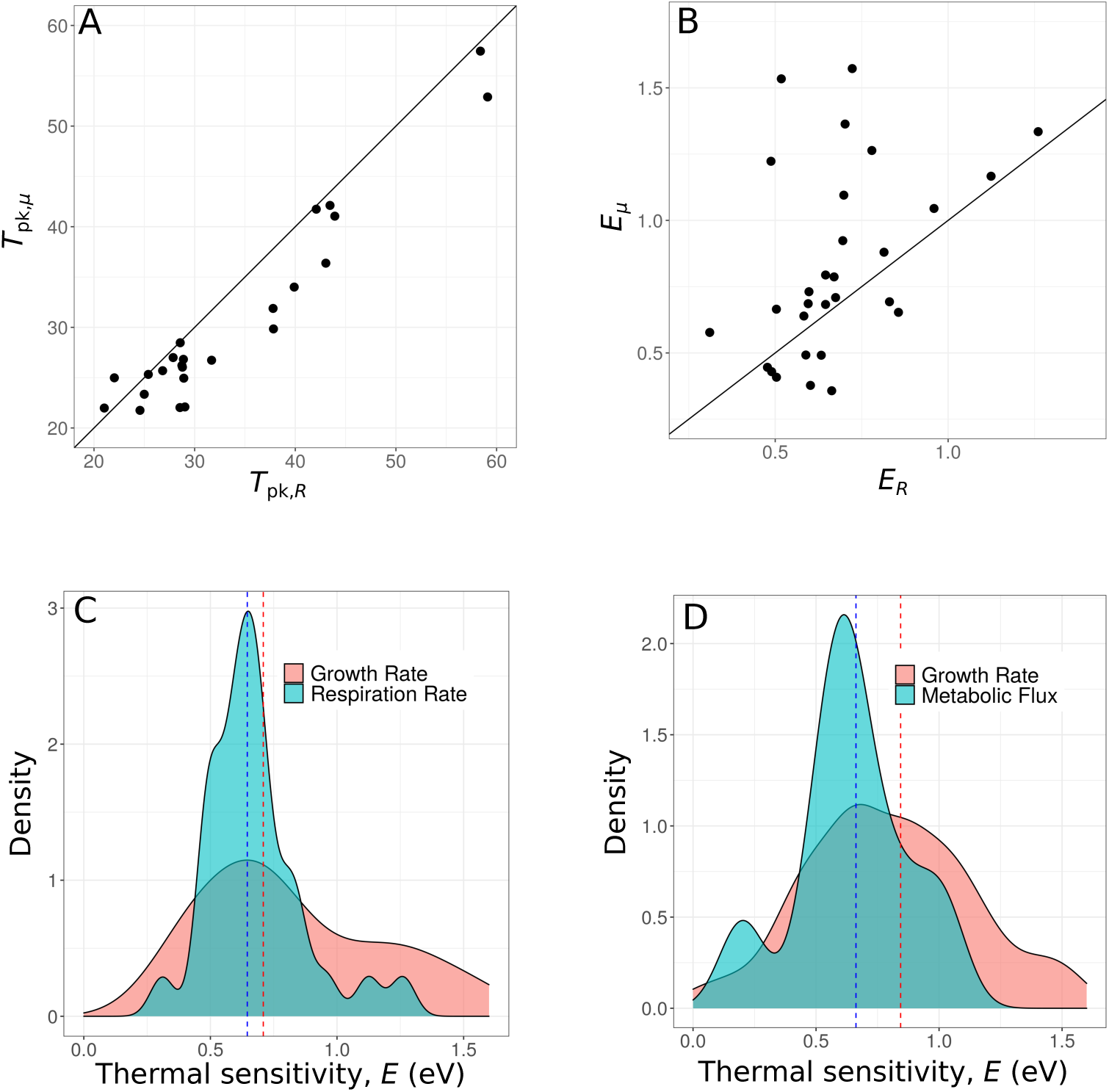
Variation in TPC parameters. **A** and **B** The relationship between growth rate and respiration rate for *T*_pk_ and thermal sensitivity (*E*) respectively (1:1 lines shown). Datapoints are parameter estimates extracted from fits to empirical data (n = 29). There are five fewer points for the *T*_pk_ comparison because one or both of the TPCs did not peak within the range of the data and thus *T*_pk_ could not be compared (however the thermal sensitivity, *E*, can still be estimated for these). **C** Distribution of *E* for growth rate and respiration rate in the experimental data. Red (growth rate) and blue (respiration rate) dotted lines show median values (median *E*_*µ*_ = 0.71, median *E*_*R*_ = 0.65). **D** Distribution of *E* for growth rate and metabolic flux rates (proxies for respiration) from a data-synthesis of > 400 bacterial TPCs ^23^ (median *E*_*µ*_ = 0.84, median *E*_*R*_ = 0.66). Median *E* values in our experimentally-derived TPCs are lower then those in the data-synthesis because the former were estimated by fitting the Boltzmann-Arrhenius model and the latter using the Sharpe-Schoolfield model (Methods).

The expectation for a decreasing CUE response to temperature is based on the assumption that respiration is more sensitive to temperature (higher *E*) than growth. However, given our theoretical analysis, our empirical results imply higher sensitivity for growth in most cases (*i.e*., *E*_*µ*_ > *E*_*R*_; Fig 1). We investigated this further using our paired growth and respiration rate TPC data. Comparing the *E*_*R*_ and *E*_*µ*_ values across strains, we find that whilst the two are positively correlated (Pearson’s *r* = 0.432), on average, *E*_*µ*_ is significantly greater than *E*_*R*_ (paired *t*_28_ = 2.513, *p* = 0.009, Fig. 3B). To determine the generality of our results, we next expanded our investigation of the difference between *E*_*µ*_ and *E*_*R*_ using a synthesis of published data spanning a much wider diversity of bacteria ^23^. We find strikingly similar differences in the shape of the distributions of *E*_*µ*_ and *E*_*R*_ in our experimental (Fig. 3C) and literature data (Fig. 3D), and find the same pattern of *E*_*µ*_ > *E*_*R*_ on average within the data-synthesis TPCs (median *E*_*µ*_ = 0.84, median *E*_*R*_ = 0.66, Fig. 3D). Therefore, the CUE TPC is more likely to increase or be thermally insensitive, than decrease within the OTR across bacteria in general.

Our results yield a new understanding of the temperature dependence of microbial carbon use efficiency. Our study on 29 strains of environmentally isolated aerobic bacteria combined with our data-synthesis goes far beyond the scope of any previous culture-based studies into the temperature dependence of CUE and its underlying traits. We find that CUE typically responds either positively to temperature, or is invariant with temperature within the OTR (Fig. 2). Focusing on the OTR of each strain is key here, as this is the temperature range within which the population typically operates, and only in the case of extreme warming events would the CUE response beyond the OTR be relevant. This general pattern in the CUE temperature dependence arises due to growth rate being typically more thermally sensitive than respiration rate (*E*_*µ*_ > *E*_*R*_, Fig. 3B). Therefore, contrary to previous thinking, we conclude that bacterial CUE generally increases or is invariant with temperature within strain-specific physiologically and ecologically meaningful temperature ranges.

The fact that growth rates generally peak at lower temperatures than respiration rates (*T*_pk,*µ*_ < *T*_pk,*R*_, Fig. 3A) is in agreement with previous empirical studies at community ^24^, as well as strain levels across both aerobic ^25,26^ and anaerobic ^27^ bacteria. Although the mechanistic basis for this systematic pattern is unclear, recent work suggests that it may be driven by stronger thermal constraints acting on carbon uptake and allocation rates, relative to respiration ^1,28^. There is very little evidence in previous studies for the differences in thermal sensitivity (*E*) of growth and respiration that we report here. Community respiration rates have in fact been reported to display higher thermal sensitivity than growth rate in aquatic systems ^14,18^. However, it is difficult to disentangle the effect of nutrient limitation on growth versus respiration in community-level measurements of these rates, and it has been suggested that growth may be more nutrient limited than respiration in such settings ^5^. Yet, a greater sensitivity of respiration relative to growth (and therefore, a negative response of CUE to temperature) is the assumption in most soil organic matter decomposition models ^3,4,15^. Indeed, it is unclear why the growth and respiration responses should display differences in thermal sensitivity without the effects of nutrient limitation. If metabolic rate is a temperature dependent process, and biomass production is fueled by metabolism, it should follow that the temperature sensitivities of each should match ^29^. However, although the responses of growth and respiration to temperature can be modelled on the basis of their responses being similar to a single rate-limiting enzymatic reaction ^30^, these rates are in reality the end result of numerous complex biochemical and physiological processes, each with their own independent thermal sensitivities ^14,31^. For aerobic heterotrophs, we may consider respiration rate to be equivalent to their “metabolic rate”, a process fundamentally dependent upon temperature ^31^. Growth (or biomass production) however is a more emergent trait based on the fraction of metabolism allocated to it ^31^. Given that the efficiency of allocation of carbon to growth varies with temperature in autotrophs ^28^, a similar constraint may exist in heterotrophs.

We found a narrower distribution of *E*_*R*_ values than *E*_*µ*_ (Fig. 3C). The differences in the shape of the distributions of *E*_*R*_ and *E*_*µ*_ in our empirical results were also reflected in the global data-synthesis (Fig. 3D), implying that this phenomenon may be generalisable across the full taxonomic diversity of bacteria. The greater variability of *E*_*µ*_ relative to *E*_*R*_ indicates that the generally positive CUE thermal response is partly due to the ability of bacterial populations to modify their carbon uptake rate and allocation efficiency for a given, constrained respiration rate. This also indicates stronger evolutionary as well as acclimation constraints acting upon the thermal sensitivity of respiration (the more fundamental metabolic process) than growth rate (the more emergent process), which can take a wider range of values. Indeed, recent work has shown that *E*_*µ*_ can escape biophysical constraints and adapt to environmental conditions ^32,33^.

Theoretical calculations have placed maximum bacterial CUE at about 0.6 ^34,35^, and similar values have been reported from pure culture experiments (CUE = 0.6-0.85) ^2,36,37^. A recent metabolic modelling study predicts variation in maximum CUE between taxa, with a range of 0.22 to 0.98 across different bacteria, with an average of ∼0.62 ^38^. This theoretical variation is realised in the wide range of bacterial CUE values obtained from isolate experiments ^39^. Generally lower CUE values are reported in natural systems than from isolate experiments ^39^, from as low as 0.01 in the most dilute systems, to nearer 0.5 in eutrophic systems ^10^. The experimental data shown here fall very much within these ranges; for all recorded measurements across all experimental conditions, median CUE = 0.22, for only the maximal CUE values recorded from each strain, median CUE = 0.38 (see Supplementary Figure S2). We see a taxonomic divergence in these CUE measurements between the two main phyla in our empirical dataset, with *Firmicutes* tending towards high efficiency whilst *Proteobacteria* are generally less efficient (Figs. 2 and S2). This is in agreement with recent work which suggests that closely related strains may have more similar CUE thermal responses than expected by chance (*i.e. E*_CUE_ is phylogenetically heritable) ^13^. Furthermore, Pold *et al*. ^13^ *show that the Q*_10_ of CUE is higher for more efficient taxa which is analogous to our result of a negative relationship between *E*_CUE_ and maximum CUE. We find that this trade-off is well described as a linear relationship, with highly negative *E*_CUE_ values not being found, suggesting a potential biological limit. Our results have extended this understanding through a more precise estimation and generalisation of variation in *E*_CUE_, via increased temperature measurements across strains adapted to a wider range of temperatures (*T*_pk,*µ*_ 22°C - 57°C).

The overall magnitude of these CUE values are likely to be an over-estimate compared to the “real” growth efficiency calculated as the total carbon uptake allocated to growth. This is due to the implicit assumption of the commonly used CUE measure (Eqn. 2) that all carbon is allocated to either growth or respiration. In reality, there may be other avenues of carbon loss that are not visible to this experiment, such as excretion of metabolites. Whether this would cause a significant difference to these results of temperature dependent CUE would depend on whether excretion displays a pattern of temperature sensitivity distinct from respiration. The release of carbon by excretion is commonly assumed to be insignificant in models of bacterial growth ^40^, however bacteria do excrete or leak metabolic by-products into the culture medium ^1,40,41^. In particular, with high levels of excess carbon in the substrate, some heterotrophic bacteria will excrete partially oxidised carbon into the environment in order to drain reducing power ^42^. When nitrogen or phosphorous are the limiting nutrients and carbon levels are high, carbon excretion levels are high ^43^. When carbon is the limiting nutrient however, levels of carbon excretion are much lower — Dauner *et al*. ^43^ *report in the region of 3-6% of carbon uptake for B. subtilis*. Our experimental data were derived from growth in the LB medium. This is a rich medium designed for exponential growth under essentially nutrient-unlimited conditions. This was used to avoid the limitations of studies from natural systems, where nutrient limitation is likely to play a major role in the CUE response ^5^. The most likely nutrient limiting growth in LB however is carbon ^44^ and therefore excretion is expected to account for only a small percentage of carbon loss. The results shown here are thus a reliable quantification of the temperature dependence of CUE in the absence of nutrient limitation.

Despite our empirical data being derived from lab experiments under nutrient saturated conditions, they represent a wide variety of strains isolated from environmental soil samples grown in a complex culture medium. Furthermore, we have extended these results to a data-synthesis spanning the entire taxonomic diversity of bacteria for which TPC data are available. Thus, our results are more generalisable, and applicable to real-world scenarios than previous culture-based experiments, which have tended to use lab-adapted strains grown on single carbon substrates, *e.g*. glucose. Our data are derived from cultures in exponential growth and therefore may provide a poor comparison to natural environments. These systems are often assumed to be at steady state, where CUE may be driven by maintenance metabolism of much lower turnover populations more generally. However, microbial systems may be more dynamic in nature, with repeated successional changes following environmental pertubations ^45^. Furthermore, environments contain ‘hot-spots’ of microbial activity with much higher process rates than average conditions ^46^, where exponential growth is relevant.

In conclusion, we have shown that, in contrast to current thinking, the response of bacterial CUE to temperature is generally invariant or positive within a biologically and ecologically relevant temperature range. This suggests that bacterial taxa are more robust to temperature change than is currently thought. These findings are important both, for physiologists aiming to understand abiotic effects on bacterial growth efficiency, as well as for parameterising ecosystem models for environment-driven variation in microbial carbon sequestration and efflux. In particular, re-parameterising microbial CUE in ecosystem models as an insensitive or increasing rather than decreasing function of temperature will likely have a major effect on predictions for both short-term responses of microbial community fluxes to temperature fluctuations, as well as longer term responses to climate change).

## Methods

### Quantifying the temperature-dependence of CUE theoretically

Here we make precise the relationship between the temperature-dependence of CUE and that of its underlying metabolic traits using a mathematical model. Consider a general equation for microbial population growth:

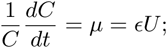

where the change in population biomass, *C*, over time, *t*, (the growth rate, *µ*) is determined by the product of the carbon uptake rate, *U*, and an efficiency, *ϵ*. This is the nutrient unlimited version of a more general growth equation appropriate for measurement of exponential population growth ^28,47^. Although there may be other sources of carbon loss to a growing bacterial population such as metabolite excretion, we assume that the majority of carbon uptake is allocated to growth and respiration, *i.e. U* ≈ *R* + *µ*. Then, *ϵ* can be expressed as:

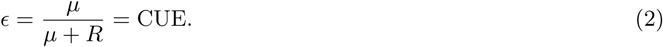

This is the same CUE (carbon use efficiency) measure found throughout the bacterial literature ^1,2,4,11,35^ (eq. 1), but this simple derivation makes explicit that the measure is meaningful only in the exponential growth phase of a population: it is (approximately) the proportion of carbon taken up by the cell that is allocated to growth during the exponential growth phase of the population.

Next, we consider how the TPC of CUE depends on TPCs of the underlying growth and respiration rates. The TPCs of a metabolic rate (*B*) can be adequately modelled by using a simplified Sharpe-Schoolfield equation ^30^ obtained by dropping the low temperature inactivation and re-expressing the equation with *T*_pk_ as an explicit parameter ^23,33,48,49^:

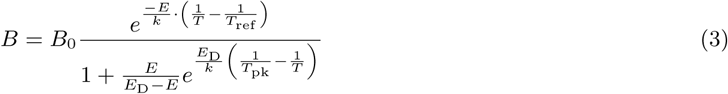

Here, *T* is temperature in Kelvin (K), *B* is a biological rate, *B*_0_ is the rate at a low reference temperature (*T*_ref_), *E* is the activation energy (eV), *E*_D_ the deactivation energy that determines the rate of decline in the biological rate beyond the temperature of peak rate (*T*_pk_), and *k* is the Boltzmann constant (8.617 × 10^*−*5^ eV K^*−*1^). The temperature-independent constant *B*_0_ includes the scaling effect of cell size, which we ignore here as cell size variation is not relevant for understanding the shape of the TPC of CUE (assuming cell size does not change significantly in the timescale over which CUE is measured). Substituting the full TPCs of *µ* and *R* defined using eq. 3 into eq. 2 can be used to quantify the CUE TPC, and can result in a large array of shapes depending upon the parameters of the *µ* and *R* TPCs (Supplementary Figure S3). However, the entire range of temperatures spanned by the TPCs of *µ* and *R* in eq. 3 are not biologically relevant because organisms generally live within their “Operational Temperature Range”(OTR), defined as the temperature range from some lower critical temperature (*e.g*., 0°C) and the temperature of peak fitness *µ* (henceforth denoted by *T*_pk,*µ*_) ^50,51^ (the “Phase 1” range in Fig. 1A). Additional phases of the TPCs of *µ, R* and CUE can also be identified — the range between the temperature of peak *µ* and peak *R* and that beyond the peak of *R* (Phase 2 and 3 respectively in Fig. 1A) — but these are also not relevant here. Within this OTR the TPCs of *µ* and *R* can be modelled simply using the Boltzmann-Arrhenius function ^23,30,51,52^, eq 4 (the numerator of eq. 2):

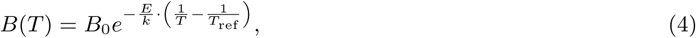

This assumes that neither growth nor respiration peak within the OTR. Indeed, growth cannot peak within the OTR by definition, as this is the range from the minimum growth temperature up to the peak growth temperature ^51^. Therefore to use the Boltzmann-Arrhenius function here, we must also assume that respiration generally peaks at higher temperatures than growth, as has previously been suggested ^1,24^. This expectation is observed within our dataset of empirical TPCs (see supplementary information). Therefore within the OTR (the typically-experienced temperature range for a strain), we can define an expression for CUE by using Boltzmann-Arrhenius functions (eq. 4) for growth (*µ*) and respiration (*R*) respectively, to give:

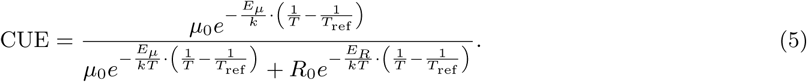

The simplification of equation 5 yields a CUE function which is monotonic over the OTR, with a direction defined entirely by differences in *E*_*µ*_ and *E*_*R*_. If *E*_*µ*_ > *E*_*R*_, CUE rises with temperature over the OTR, if *E*_*µ*_ < *E*_*R*_, CUE declines with temperature across the OTR. This is the basis for previous theoretical expectations for the CUE temperature response ^1^, here formalised as eq. 5. Specifically, we can approximate the denominator in eq. 5 using a Taylor series expansion, to obtain the following approximation for CUE:

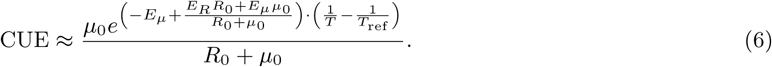

This equation has the form of a Boltzmann-Arrhenius function with:

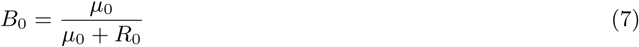

and the apparent activation energy (a measure of thermal sensitivity of CUE) as

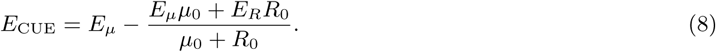

Thus, the CUE TPC is necessarily monotonic within the OTR as long as *T*_pk,*µ*_ < *T*_pk,*R*_ (as is almost always the case; see Fig. 3).

This expression can be used to determine the direction of the CUE thermal sensitivity within the OTR as follows. Recognising that the condition for CUE to decrease with temperature is *E*_CUE_ < 0, we can rearrange eq 8 as :

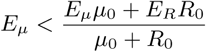

This simplifies to the condition

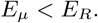

That is, within the OTR, CUE increases if *E*_*R*_ < *E*_*µ*_ (⇒ *E*_CUE_ > 0), decreases if *E*_*R*_ > *E*_*µ*_ (⇒ *E*_CUE_ < 0), and is insensitive to temperature if *E*_*R*_ = *E*_*µ*_(⇒ *E*_CUE_ = 0)

### Quantifying the temperature dependence of CUE experimentally

We used 29 strains of environmentally isolated aerobic bacteria from our laboratory culture collection (see supplementary table S1). These strains were isolated under a range of different temperatures for a species sorting experiment, aiming to reconstruct the wide diversity of bacterial temperature fitness present in soils. We experimentally quantified the TPC of CUE for these bacteria as follows.

At each experimental temperature, frozen bacterial cultures were revived and grown to carrying capacity at the experimental temperature (acclimation period - to restrict influence of temperature stress on TPC, or equalise it across experimental points). Revived cultures were grown in LB medium in replicates of 4 and growth rate and respiration rate were measured during exponential growth using flow cytometry cell counts (growth) and MicroResp™ (respiration). This was repeated across a range of temperatures spanning the full TPC for each isolate.

From the flow cytometry measurements, estimates of carbon biomass in the cultures were made based on cell diameters ^53^, and growth in the exponential phase calculated as:

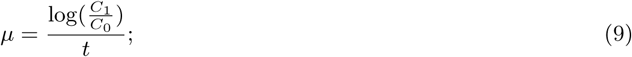

where *C*_0_ is the starting biomass, *C*_1_ is the final biomass and *t* is the duration of the experiment. MicroResp™ was used to give a quantitative measure of the cumulative respired CO_2_ produced during the growth experiment ^54^. From this, the per-capita respiration rate was calculated in terms of carbon mass, according to:

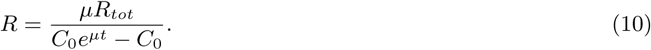

Here, *R*_*tot*_ is the total mass of carbon produced, *C*_0_ is the initial population biomass, *µ* is the previously calculated growth rate and *t* is the duration of the experiment (see supplementary material for full details of the derivation of eq 10). This measure of respiration rate is directly comparable to the specific growth rate, *µ*, and overcomes a problem shared by practically all previous empirical measurements of CUE. Specifically, for a given temperature, previous methods have often required growth rates to be measured at a different timescale, or at a different time point of population growth, than the measurement of respiration rate. This is because *µ* needs to be measured over time-period sufficiently long enough to allow changes in cell density to be detectable using optical methods, while respiration rate can be measured over much shorter timescales. The resulting difference in timescales of measurement permits a greater level of thermal acclimation of growth relative to respiration. Furthermore, in cases where these measurements *have* been made over a similar time-frame, respiration rates are often normalised only to the starting mass of the growing population, and neglect to include changes in the growing population size over time (*e.g*. Keiblinger *et al*. ^11^, *Créach et al*. ^55^, *Warkentin et al*. ^56^). Indeed, direct comparisons of the TPCs of growth and respiration that our methods allow have largely been lacking from the literature, making it difficult to link these processes to temperature-dependent CUE at the appropriate timescale ^57^.

#### Calculating CUE from the experimental data

Having measured the TPCs of growth and respiration rate, we then calculated the within-OTR TPC of CUE for each bacterial strain as follows. We first fit the Sharpe-Schoolfield model (eq. 3, Methods) to paired growth rate and respiration rate TPCs for each of the 29 strains of aerobic bacteria to determine the respective *T*_pk,*µ*_ and *T*_pk,*R*_, and then fitted the Boltzmann-Arrhenius model (eq. 4) to the TPC from the rate at minimum temperature up to its *T*_pk_. To fit eq. 4 to the temperature dependent growth and respiration rates to each of the 29 strains in our dataset, we used only those strains that had at least 3 datapoints in the temperature range lower than their Shape-Schoolfield calculated *T*_pk_. We input these TPC parameters for *µ* and *R* (calculated from eq. 4) into eq. 5 to calculate the CUE TPC, and and its corresponding *E*_CUE_ using eq. 8. All analyses and model fitting were performed in R ^58^, using the “minpack.lm” package for non-linear least squares fitting.

#### Accounting for uncertainty in model fitting

To account for uncertainty in the estimated TPCs (*i.e*., in the parameters *B*_0_ and *E*; eq. 4) in our tests of whether the emergent CUE responds significantly to temperature, we implemented a bootstrapping approach as follows. For each strain we re-sampled the data with replacement 1,000 times and re-fit the Boltzmann-Arrhenius model (eq. 4) to the sub-sampled growth and respiration dataset. As the data are paired (each CUE value is derived from a growth and a respiration measurement), we re-sampled growth and respiration paired points (rather than re-sampling growth and respiration separately), in order to account for their covariance. From each of the paired BA model fits we calculated *E*_CUE_ according to eq. 8, obtaining a distribution of these values. We then calculated the 95% confidence interval for *E*_CUE_ as the 2.5th and 97.5th percentiles of this distribution. We asked whether or not the CIs include zero, as a robust test to determine a thermal response significantly different from a temperature insensitive response (Fig. 2).

In order to calculate a confidence envelope around each CUE TPC, we took the fitted parameters from the 1,000 bootstrapped curves for each strain and interpolated CUE curves across the temperature range for plotting. At each temperature, we took the 2.5th and 97.5th percentiles of the CUE distribution as the upper and lower bounds of the 95% confidence envelope.

### Data-synthesis of bacterial thermal performance curves

To understand our results in a broader context, we compared the thermal sensitivities of our empirically derived *µ* and *R* TPCs to those in our recent global data synthesis ^23^. This data synthesis is primarily composed of growth rate TPCs (416 bacterial *µ* TPCs), but also contains 22 bacterial metabolic flux TPCs which we use as proxies for respiration rate TPCs. This is a taxonomically and functionally diverse dataset, spanning 13 bacterial phyla and practically the entire range of thermal niches inhabited by bacteria. Rather than re-analyse the raw data here, we directly take the *E*_*µ*_ and *E*_*R*_ estimates provided and compare the distributions to those of our empirically derived TPCs. The data-synthesis calculates *E* directly from the Sharpe-Schoolfield model (eq. 3), whereas here we calculate *E* from the Boltzmann-Arrhenius function (eq. 4) fitted within the OTR. This is expected to cause a difference in the overall magnitude of *E* between datasets (lower *E* using Boltzmann-Arrhenius due to curvature as trait values approach *T*_pk_ ^51^), however we emphasise this does not affect *E*_*µ*_ and *E*_*R*_ comparisons within these datasets, nor the comparison of distributions between these datasets.

## Acknowledgements

TPS was supported by a BBSRC DTP scholarship (BB/J014575/1). TB and SP were funded by NERC grants NE/M020843/1 and NE/S000348/1.

