## Supplementary Information for "Systematic variation in the temperature dependence of bacterial carbon use efficiency"

#### Contents

**1 Derivation of biomass-specific respiration rate equation 1**

**2 Strains used for empirical CUE measurements 2**

**3 Testing $E_{\mathbf{CUE}}$ directionality using bootstrapped confidence intervals 4**

**4 Distribution of CUE values 6**

**5 CUE thermal responses modelled with Sharpe-Schoolfield curves 6**

#### 1 Derivation of biomass-specific respiration rate equation

The MicroResp system provides a measure of the total cumulative $CO_2$ produced over the duration of incubation and has been used to quantify bacterial respiration[1,2,3]. Assuming that respiration is a mass-specific rate that is constant across the experiment and that $CO_2$ accumulates under the assumption of negligible escape or re-absorption, the rate of change of cumulative total $CO_2$ respired (i.e. captured by the MicroResp system) ($R_{\text{tot}}$) can be written:

$$\frac{dR_{\text{tot}}}{dt} = RC(t); \tag{1}$$

where the change in total respired carbon ($R_{\text{tot}}$) at time $t$ is given by the product of the mass-specific rate of respiration $R$ and the population biomass at time $t$, $C(t)$, i.e. the rate of change of accumulated $CO_2$ is proportional to the population size. Given that the cultures are in exponential growth, the population size at any time point is given by:

$$C(t) = C_0 e^{\mu t}; \tag{2}$$

where $C_0$ is the initial population size. Substituting Eq. 2 into Eq. 1 gives:

$$\frac{dR_{\text{tot}}}{dt} = RC_0 e^{\mu t}. \tag{3}$$

We can integrate this expression across time to get the predicted total respiration at any time point. This gives:

$$R_{tot}(t) = C_1 + \frac{C_0 R e^{\mu t}}{\mu}, \tag{4}$$

where $C_1$ is the constant of integration. Setting $t = 0$ and solving for $C_1$ we get:

$$R_{\text{tot}}(t) = R_{\text{tot}}(0) + \frac{C_0 R e^{\mu t}}{\mu} - \frac{C_0 R}{\mu}, \tag{5}$$

which we can re-arrange to give $R$, the biomass specific respiration rate:

$$R = \frac{\mu(R_{\text{tot}}(t) - R_{\text{tot}}(0))}{C_0 e^{\mu t} - C_0}. \tag{6}$$

Given that the cumulative respired $CO_2$ at $t(0) = 0$, we can simplify this in the form of eq. 7 (main text Eq. 10):

$$R = \frac{\mu R_{\text{tot}}}{C_0 e^{\mu t} - C_0}. \tag{7}$$

#### 2 Strains used for empirical CUE measurements

We used 29 strains of environmentally isolated aerobic bacteria from our laboratory culture collection, adapted to a wide range of thermal niches. Full details of all the strain used and their taxonomy as determined through 16S sequencing is shown in Table S1.

Table S1: **List of strains used for empirical CUE measurements.**

Strain names refer to internal ID codes. Taxonomy to genus level determined via 16S sequences. Peak growth temperature ($T_{\mathrm{pk},\mu}$), estimated through Sharpe-Schoolfield model fits, to show the differing thermal niches between strains. * = unreliable $T_{\mathrm{pk},\mu}$ estimate from model fitting due to lack of datapoints beyond the peak; $T_{\mathrm{pk},\mu}$ estimated by eye.

| Strain | Phylum | Class | Order | Family | Genus | $T_{\mathrm{pk},\mu}$ (°C) |
|---|---|---|---|---|---|---|
| 04_RT_02 | Proteobacteria | Gammaproteobacteria | Pseudomonadales | Pseudomonadaceae | Pseudomonas | 23.35 |
| 04_RT_05 | Proteobacteria | Gammaproteobacteria | Pseudomonadales | Pseudomonadaceae | Pseudomonas | 27.00 |
| 04_04_02 | Proteobacteria | Gammaproteobacteria | Pseudomonadales | Pseudomonadaceae | Pseudomonas | 26.82 |
| 04_04_04 | Proteobacteria | Betaproteobacteria | Burkholderiales | Oxalobacteraceae | Collimonas | 21.75 |
| 10_RT_01 | Proteobacteria | Gammaproteobacteria | Pseudomonadales | Pseudomonadaceae | Pseudomonas | 26.57 |
| 10_RT_02 | Proteobacteria | Gammaproteobacteria | Pseudomonadales | Pseudomonadaceae | Pseudomonas | 25.69 |
| 10_10_06 | Actinobacteria | Actinobacteria | Micrococcales | Micrococcaceae | Arthrobacter | 22.09 |
| 21_RT_05 | Actinobacteria | Actinobacteria | Corynebacteriales | Nocardiaceae | Nocardia | 26.72 |
| 21_21_02 | Proteobacteria | Gammaproteobacteria | Pseudomonadales | Pseudomonadaceae | Pseudomonas | 24.98 |
| 21_21_04 | Proteobacteria | Betaproteobacteria | Burkholderiales | Oxalobacteraceae | Collimonas | 22.03 |
| 21_21_06 | Proteobacteria | Gammaproteobacteria | Pseudomonadales | Pseudomonadaceae | Pseudomonas | 21.98 |
| 30_RT_01 | Proteobacteria | Betaproteobacteria | Burkholderiales | Comamonadaceae | Variovorax | 24.95 |
| 40_RT_01 | Firmicutes | Bacilli | Bacillales | Bacillaceae | Bacillus | 36.38 |
| 40_RT_03 | Firmicutes | Bacilli | Bacillales | Planococcaceae | Viridibacillus | 27.03 |
| 40_RT_04 | Firmicutes | Bacilli | Bacillales | Planococcaceae | Viridibacillus | 26.73 |
| 50_RT_01 | Firmicutes | Bacilli | Bacillales | Bacillaceae | Bacillus | 35.71 |
| 04_04_06 | Proteobacteria | Gammaproteobacteria | Pseudomonadales | Pseudomonadaceae | Pseudomonas | 28.47 |
| 30_RT_02 | Proteobacteria | Betaproteobacteria | Burkholderiales | Comamonadaceae | Variovorax | 25.32 |
| 21_RT_03 | Firmicutes | Bacilli | Bacillales | Paenibacillaceae | Paenibacillus | 26.21 |
| 30_30_08 | Proteobacteria | Alphaproteobacteria | Rhizobiales | Xanthobacteraceae | Labrys | 26.04 |
| 40_RT_05 | Firmicutes | Bacilli | Bacillales | Bacillaceae | Bacillus | 29.84 |
| 40_40_01 | Firmicutes | Bacilli | Bacillales | Bacillaceae | Bacillus | 31.88 |
| 40_40_03 | Firmicutes | Bacilli | Bacillales | Bacillaceae | Bacillus | 34.00 |
| 40_40_04 | Firmicutes | Bacilli | Bacillales | Planococcaceae | Rummeliibacillus | 41.74 |
| 40_40_05 | Firmicutes | Bacilli | Bacillales | Paenibacillaceae | Cohnella | 42.12 |
| 40_40_06 | Firmicutes | Bacilli | Bacillales | Planococcaceae | Viridibacillus | 41.05 |
| 50_50_01 | Firmicutes | Bacilli | Bacillales | Planococcaceae | Rummeliibacillus | 52.89 |
| 50_50_03 | Firmicutes | Bacilli | Bacillales | Paenibacillaceae | Brevibacillus | 57.45 |
| 50_50_04 | Firmicutes | Bacilli | Bacillales | Paenibacillaceae | Brevibacillus | 55* |

### 3 Testing $E_{\mathrm{CUE}}$ directionality using bootstrapped confidence intervals

To provide a statistical test for the directionality of the CUE thermal response, *i.e.* whether it differs signficantly from zero, we used a bootstrapping approach. For each strain we re-sampled the data with replacement 1,000 times and re-fit the Boltzmann-Arrhenius model (eq. 4, main text) to the sub-sampled growth and respiration dataset. From each of the paired BA model fits we calculated $E_{\mathrm{CUE}}$ according to eq. 8 (main text), obtaining a distribution of values. We then calculated the 95% confidence interval for $E_{\mathrm{CUE}}$ as the 2.5th and 97.5th percentiles of this distribution. To test the directionality, we finally asked whether or not the CIs include zero. For most strains the distribution of bootstrapped $E_{\mathrm{CUE}}$ is fairly normal, however there are some instances of high skewing due to an imbalanced number of datapoints across temperatures within the OTR (Fig. S1). This results in asymmetric CIs (main text Fig. 2A). Full results of the bootstrapped directionality test shown in table S2.

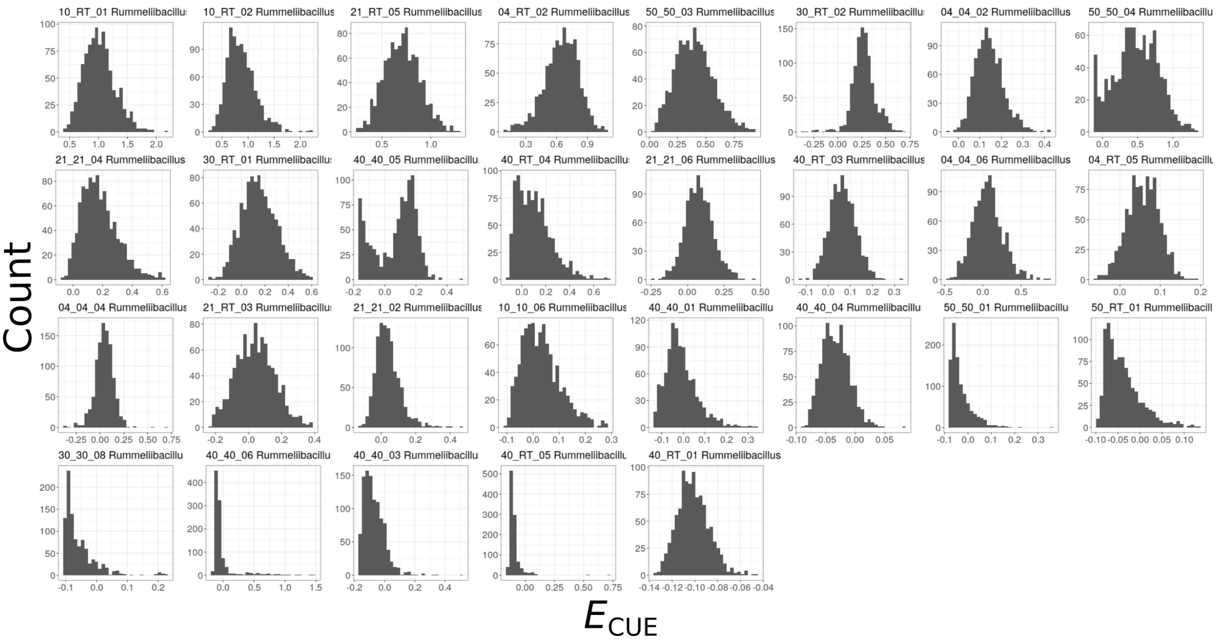


Figure S1: **Distribution of bootstrapped $E_{\mathrm{CUE}}$.**

Table S2: **Bootstrapped CUE thermal response results.**

Results of tests for the directionality of the CUE thermal response for each strain. $E_{\mathrm{CUE}}$ min and $E_{\mathrm{CUE}}$ max refer to the lower and upper 95% CIs for $E_{\mathrm{CUE}}$, obtained by bootstrapping. If the CIs fall above zero, CUE increases with temperature; if the CIs fall below zero, CUE decreases with temperature; if the CIs include zero, CUE is statistically invariant with temperature.

| Strain | $E_\mu$ | $E_R$ | $E_{\mathrm{CUE}}$ median | $E_{\mathrm{CUE}}$ min | $E_{\mathrm{CUE}}$ max | Result |
|---|---|---|---|---|---|---|
| 10_RT_01 | 1.52 | 0.52 | 0.99 | 0.57 | 1.59 | Increasing |
| 10_RT_02 | 1.56 | 0.72 | 0.83 | 0.43 | 1.51 | Increasing |
| 21_RT_05 | 1.22 | 0.49 | 0.71 | 0.38 | 1.08 | Increasing |
| 04_RT_02 | 1.37 | 0.70 | 0.66 | 0.27 | 0.95 | Increasing |
| 50_50_03 | 1.10 | 0.70 | 0.39 | 0.11 | 0.75 | Increasing |
| 30_RT_02 | 0.58 | 0.31 | 0.27 | 0.04 | 0.52 | Increasing |
| 04_04_02 | 0.79 | 0.65 | 0.14 | 0.03 | 0.29 | Increasing |
| 50_50_04 | 1.26 | 0.78 | 0.47 | -0.09 | 1.06 | Invariant |
| 21_21_04 | 0.93 | 0.70 | 0.16 | -0.01 | 0.47 | Invariant |
| 30_RT_01 | 0.65 | 0.50 | 0.14 | -0.11 | 0.46 | Invariant |
| 40_40_05 | 0.72 | 0.60 | 0.11 | -0.16 | 0.26 | Invariant |
| 40_RT_04 | 0.78 | 0.67 | 0.09 | -0.08 | 0.45 | Invariant |
| 21_21_06 | 1.04 | 0.95 | 0.08 | -0.10 | 0.28 | Invariant |
| 40_RT_03 | 0.69 | 0.60 | 0.06 | -0.05 | 0.18 | Invariant |
| 04_04_06 | 1.32 | 1.27 | 0.06 | -0.29 | 0.50 | Invariant |
| 04_RT_05 | 0.88 | 0.81 | 0.06 | -0.02 | 0.13 | Invariant |
| 04_04_04 | 0.64 | 0.58 | 0.05 | -0.15 | 0.22 | Invariant |
| 21_RT_03 | 1.17 | 1.13 | 0.04 | -0.18 | 0.28 | Invariant |
| 21_21_02 | 0.71 | 0.67 | 0.03 | -0.07 | 0.21 | Invariant |
| 10_10_06 | 0.69 | 0.65 | 0.02 | -0.08 | 0.20 | Invariant |
| 40_40_01 | 0.45 | 0.48 | -0.02 | -0.12 | 0.17 | Invariant |
| 40_40_04 | 0.43 | 0.49 | -0.03 | -0.07 | 0.01 | Invariant |
| 50_50_01 | 0.50 | 0.59 | -0.05 | -0.07 | 0.08 | Invariant |
| 50_RT_01 | 0.40 | 0.51 | -0.05 | -0.09 | 0.07 | Invariant |
| 30_30_08 | 0.65 | 0.85 | -0.07 | -0.10 | 0.07 | Invariant |
| 40_40_06 | 0.70 | 0.83 | -0.08 | -0.14 | 0.73 | Invariant |
| 40_40_03 | 0.49 | 0.63 | -0.08 | -0.16 | 0.09 | Invariant |
| 40_RT_05 | 0.36 | 0.66 | -0.11 | -0.14 | 0.03 | Invariant |
| 40_RT_01 | 0.38 | 0.60 | -0.10 | -0.13 | -0.07 | Decreasing |

#### 4 Distribution of CUE values

In figure S2, we show the distribution of calculated CUE values. Across all strains and temperature conditions (fig S2A), we find median CUE = 0.25. Splitting these by phylum, we find *Actinobacteria* (red) median CUE = 0.31, *Firmicutes* (blue) median CUE = 0.29 and *Proteobacteria* (yellow) median CUE = 0.09. Many of these values are lower than might normally be expected as they were recorded under far from optimal growth conditions (e.g. high temperature stress). Considering only the maximum CUE obtained from each strain (fig. S2B), we find *Actinobacteria* median CUE = 0.58, *Firmicutes* median CUE = 0.47 and *Proteobacteria* median CUE = 0.15, with a median across the whole dataset of 0.44. These results are more in fitting with the maximal CUE estimates across strains modelled by Saifuddin *et al.*[4].

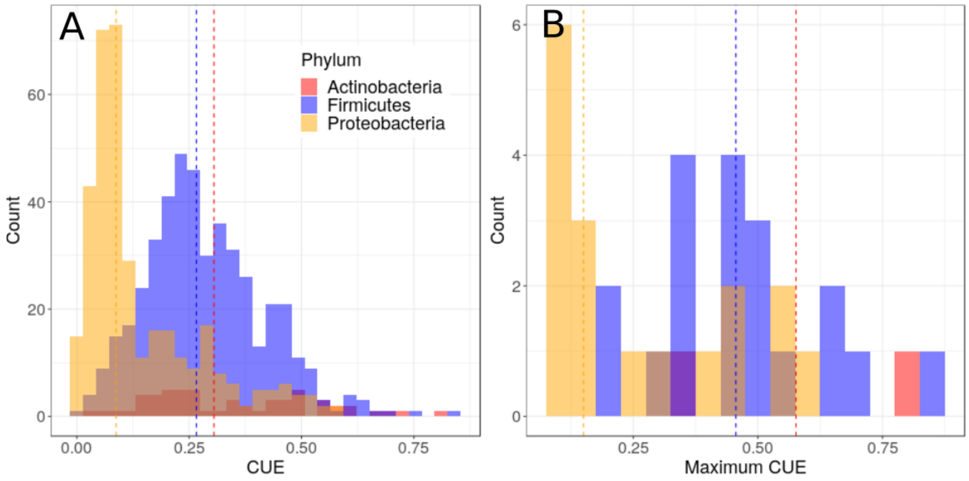


Figure S2: **Distribution of calculated CUE values. A.** Distribution of all recorded CUE values separated by phylum. Dashed lines indicate medians. **B.** Distribution of maximal CUE values recorded for each bacterial strain.

#### 5 CUE thermal responses modelled with Sharpe-Schoolfield curves

In the main text we state that Sharpe-Schoolfield curves (main text Eq. 3) defined for growth rate and respiration rate substituted into Eq. 2 (main text) could be used to investigate the thermal response of CUE. This however results in a wide array of non-linear responses, which is problematic for understanding the drivers of these responses. Examples of the shapes these curves may take across an organism's entire temperature range are given in Fig. S3, based on differences in $E$ and $T_{pk}$ between growth and respiration. We find that differences in these parameters cause the CUE curve to produce an initial increase or decrease in CUE with temperature (due to differences in E) followed by an inflection either upwards or downwards (due to differences in $T_{pk}$). The resultant CUE curves may therefore take different forms based on how consistently growth and respiration show these differences in thermal performance. In order to provide a simpler understanding of the CUE thermal response, across the temperature range normally experienced by an organism, we instead used Boltzmann-Arrhenius equations within the OTR, where differences in $E$ are the main driver and inflections due to $T_{pk}$ differences are not considered (see main text). This approach is validated by our finding that respiration peaks at higher temperatures than growth in almost all cases (main text Fig. 3)

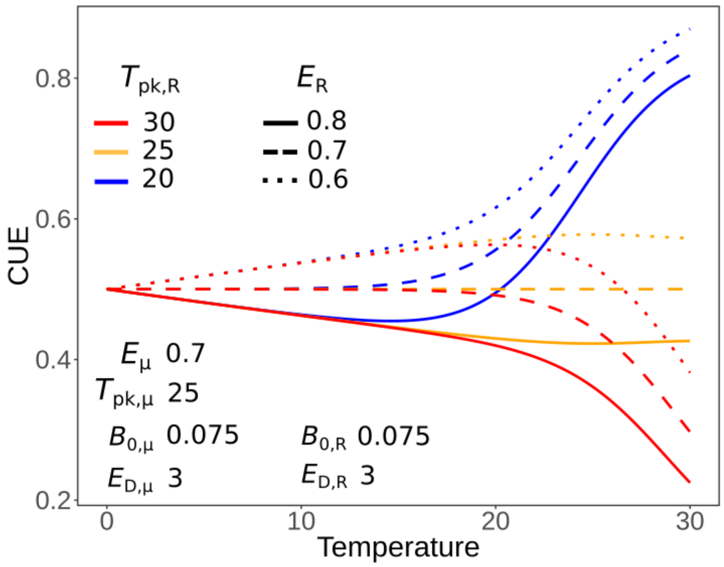


Figure S3: **Variation in CUE with temperature modelled using simulated Sharpe-Schoolfield curves.** Combining variation in $E$ and $T_{pk}$ we can see the varied shapes that a CUE TPC may take. Here, the parameters for growth rate remain constant, whilst the respiration $E$ and $T_{pk}$ parameters are allowed to vary above and below those of growth.
